# Augmentation of task-relevant variability enhances consolidation of motor learning

**DOI:** 10.1101/2021.06.03.446469

**Authors:** Mattia Pagano, Gaia Stochino, Maura Casadio, Rajiv Ranganathan

## Abstract

Motor memories undergo a period of consolidation before they become resistant to the practice of another task. Although movement variability is important in motor memory consolidation, its role is not fully understood in redundant tasks where variability can exist along two orthogonal subspaces (the ‘task space’ and the ‘null space’) that have different effects on task performance. Here, we used haptic perturbations to augment variability in these different spaces and examined their effect on motor memory consolidation. Participants learned a shuffleboard task, where they held a bimanual manipulandum and made a discrete throwing motion to slide a virtual puck towards a target. The task was redundant because the distance travelled by the puck was determined by the sum of the left and right hand speeds at the time of release. After participants initially practiced the task, we used haptic perturbations to introduce motor variability in the task space or null space, and subsequently examined consolidation of the original task on the next day. We found that regardless of the amplitude, augmenting variability in the task space resulted in significantly better consolidation. This benefit of increasing task space variability was likely due to the fact that it did not disrupt the pre-existing coordination strategy. These results suggest that the effects of variability on motor memory consolidation depend on the interplay between the induced variability and the pre-existing coordination strategy.

## Introduction

After practice at a task, motor memories undergo a period of consolidation where they transform from a ‘fragile’ state to a ‘stable’ state (Krakauer and Shadmehr 2006; Walker et al. 2003). The evidence for such consolidation can be observed because memories that are still in the fragile state can be disrupted by the practice of a related but different task, termed the ‘interfering task’. Prior work has shown the time course of this consolidation process, primarily using tasks that involve sequence learning (Robertson et al. 2004), visuomotor adaptation (Krakauer et al. 2005) and force field adaptation (Brashers-Krug et al. 1996), although there has also been some evidence against the consolidation hypothesis in certain contexts (Goedert and Willingham 2002).

A key issue in these consolidation paradigms is the nature of the interfering task. Although initial work used interfering tasks that required a ‘different’ motor response from the originally learned task - for e.g. learning a visuomotor rotation in the opposite direction (Krakauer et al. 1999) or learning a counterclockwise force field after learning a clockwise force field (Brashers-Krug et al. 1996), recent work has examined tasks that use of variations of the ‘same’ motor response by adding variability (Wymbs et al. 2016). In a series of experiments, Wymbs et al. demonstrated that adding variability to a consolidated skill could in fact further strengthen the motor skill. The study by Wymbs et al. focused primarily on the issue of ‘reconsolidation’, i.e., bringing an already consolidated memory back into a fragile state, but the role of variability during initial consolidation is still poorly understood.

Two specific issues arise when understanding the role of variability in consolidation. First, as mentioned earlier, the focus of much of the prior work has been on adaptation and sequence learning tasks, which are qualitatively different from precision tasks that require control of motor variability (e.g. a dart throw). This choice of a precision task could be especially critical considering that the interfering task involves manipulating motor variability. Second, when the task has redundancy, motor variability can be introduced along two different ‘subspaces’ – in a task space where it affects task performance, or in a null space where it has no effect on task performance (Cusumano and Cesari 2006; Latash et al. 2002; Mosier et al. 2005; Scholz and Schöner 1999; Todorov and Jordan 2002). Therefore, it is not known if the subspace in which variability is introduced has differential effects on motor memory consolidation.

The purpose of the current study was to examine the effect of introducing variability on initial motor memory consolidation. Participants practiced a precision task and motor variability was introduced through haptic perturbations in the task and null spaces (Cardis et al. 2018). We examined performance on the next day to examine how introducing variability affected retention of learning. We addressed the effects of the subspace in which variability is introduced, and amplitude of variability introduced on consolidation.

## Methods

### Participants

72 college-aged adults (aged 18-24, 25 males, 47 females, 66 right-handed) with no history of orthopaedic injuries or neurological disorders participated in the study. Each participant provided written informed consent in accordance with a protocol approved by the Michigan State University Institutional Review Board.

### Apparatus

The task was performed with a bimanual planar manipulandum (KINARM Endpoint Robot, BKIN Technologies, ON). The robot has two separate robotic arms, moving in the horizontal plane, whose end-effectors can be held by the subjects. Participants were seated on a height-adjustable chair and were instructed to look into a semi-silvered mirror screen positioned at around 45 degrees angle below eye level. Participants could not directly see their hands during the experiment, but viewed the virtual objects projected on the screen in the same plane as their hands.

### Task

Participants performed a virtual shuffleboard task that required sliding a puck toward a target (Cardis et al. 2018). The goal of the task was to slide the puck with just the right speed so that it would stop at the center of a virtual target.

At the start of each trial, to ensure a fixed starting position for both hands, participants were first instructed to move both handles to reach on the screen two home positions, close to the body. Once the home locations were reached, the individual cursors for each hand disappeared and were replaced by a puck appearing at the average position of the two hands. Participants also saw a rectangular slot positioned at distance of 10 cm ahead of the puck. Participants were then instructed to slide the puck toward this slot by making a smooth forward movement with both hands (**Figure 1A**).

**Figure 1.**
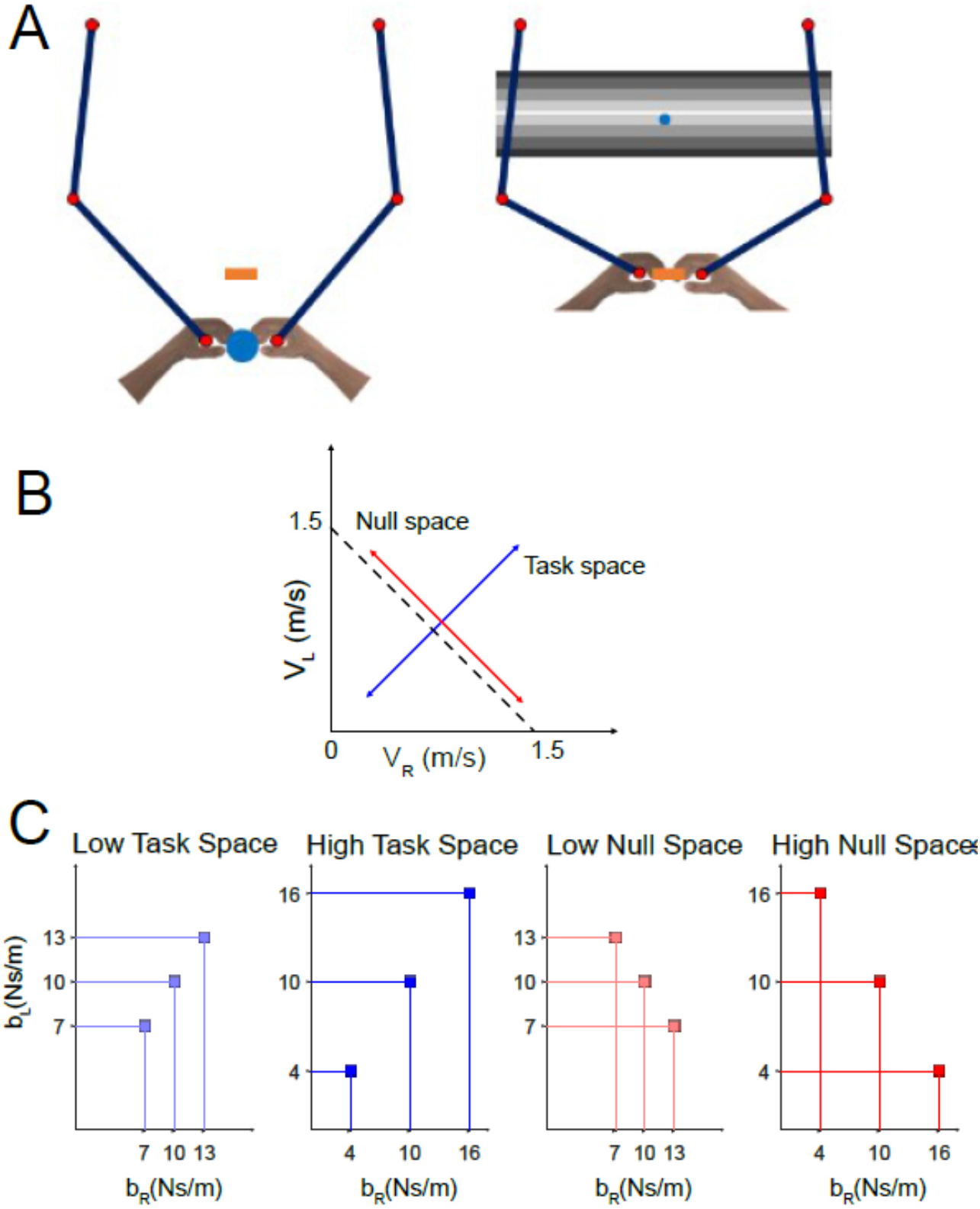
**(A) Task description.** Participants performed a virtual shuffleboard task to slide a virtual puck toward a target. Participants saw a blue puck that was shown at the midpoint of the two hands and were instructed to make a smooth forward movement with both hands toward a rectangular ‘slot’ straight ahead. Once the puck crossed the slot it was ‘released’ and participants could see it sliding on the screen toward a virtual greyscale target. **(B) Description of task and null spaces for the shuffleboard task**. The speed of the puck was determined by the sum of the left and right hand speeds at release, with perfect performance occurring when V_L_ + V_R_ = 1.5 m/s. **(C) Introducing variability in task and null spaces**. Trial to trial variability was introduced by changing the viscosity coefficient on the robot from its original value of 10 Ns/m. For introducing variability in the task space, the coefficients were positively covaried (i.e. both hands were sped up or slowed down on any given trial) whereas for the null space the coefficients were negatively covaried (i.e. one hand was sped up while the other was slowed down). The amount of variability introduced was controlled by the magnitude of the change in the viscosity coefficients.

When participants made the forward motion, the virtual puck was ‘released’ from the hands as it crossed the slot. Once released, participants were shown a screen with the virtual puck sliding toward a target (shown as a greyscale board). The final stopping location of the puck was based on the speed of the puck at release. On each trial, the motion of the puck was constrained to one-dimension (i.e. the forward direction), and was determined by both hand velocities at the instant of release as follows:

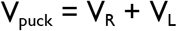

Where V_puck_ is the speed of the puck, and V_R_ and V_L_ are the right and left hand speeds (i.e. overall magnitude). To land the puck exactly on the center of the target, the puck had to be released with a speed of 1.5 m/s.

Once the puck stopped, at the end of each trial, participants were shown a score that depended on the distance of the final position of the puck from the center of the target (**Figure 1B**). This score was computed using the following equation:

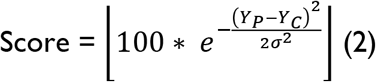

Where *Y*_*P*_ is the final position of the puck, *Y*_*C*_ is the Y position of the centre of the screen (30cm) and *σ* was empirically set equal to 3. Thus, participants achieved the highest score of 100 when the speed of the puck at release was equal to 1.5 m/s and the puck stopped exactly at the center of the target. Participants were shown the score at the end of each trial and a running average of all the scores in the trials in that block. At the end of each block of practice (50 trials), participants were provided with a mean score for that block and encouraged to improve it on the following block.

Similar to prior work (Cardis et al. 2018), velocity-dependent viscous force fields were used to provide haptic forces during the task. The force field on each hand was determined by its viscosity coefficients according to the following equation.

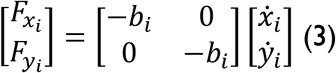

Where 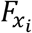 and 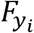 are the *x, y* components of the force field generated by the manipulandum, 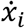 and 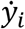 are the component of the hand velocity vector for the i-hand (i.e. i=R for the right and i=L for the left one) and *b*_*i*_ are the viscosity coefficients. The viscosity coefficients were used to manipulate the variability introduced during practice.

### Experimental Protocol

The protocol consisted of three sessions across 2 days and followed an A-B-A paradigm employed in consolidation studies (**Figure 2**). In the first and third sessions (i.e. the original task A), participants performed the shuffleboard task with no haptic perturbations. In the second session, participants performed a task (the interfering task B) that was the same as task A but with the addition of haptic perturbations from trial-to-trial to manipulate motor variability depending on which group the participants belonged to. The first two sessions were performed consecutively in the same day, with no delay between them, while the third session was performed 24 hours later. A small ‘reactivation’ block R (which consisted of 15 trials of the original task A) was introduced before the interfering task B. Although this block was not critical in the current experiment since there was no delay between practice of tasks A and B, it was introduced to keep the design consistent with prior studies where a delay was introduced between the two tasks (Wymbs et al. 2016).

**Figure 2.**
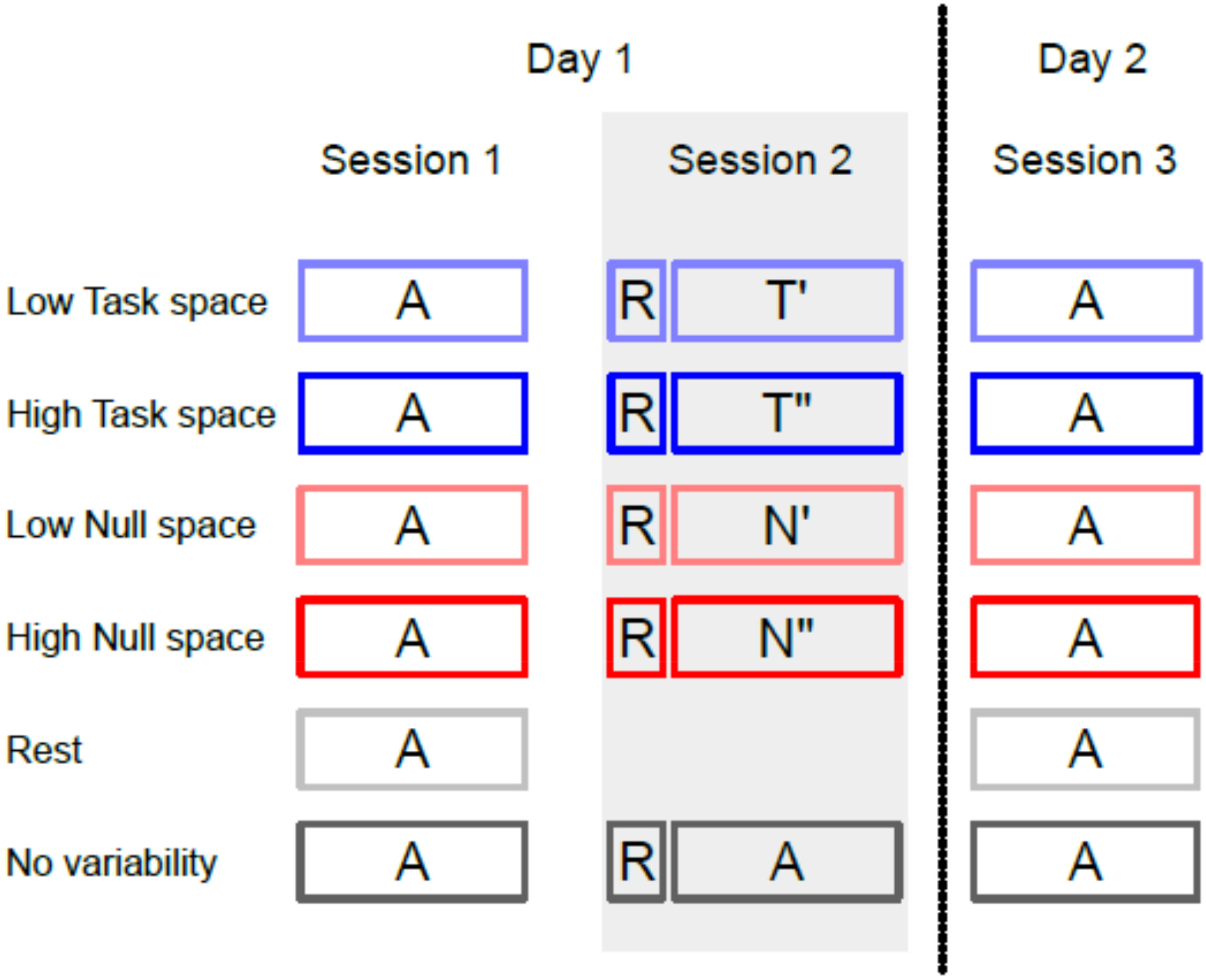
Experimental protocol. Participants performed three sessions of practice in an A-B-A paradigm. Session 1 and 2 were performed consecutively the same day, while the third session was performed 24 hours later. Participants were randomly assigned to 4 groups (n=12 participants/group) in a 2 × 2 design that varied the space in which variability was introduced (task T or null space N), and the amount of variability introduced (high or low). We also added two additional control groups of participants that either rested or performed the same task A with no variability in the second session. All participants performed the same task in sessions 1 and 3, while the task in session 2 varied depending on group membership. Consolidation was assessed by examining performance in session 3 relative to the performance at the end of session 1. A short reactivation block R (which was the same as task A) was introduced prior to task B for consistency with prior work.

### Procedure

Prior to practice of task A, all participants first performed a familiarization block of 15 trials with the robot providing no forces, to familiarize themselves with the task and the scoring system. During task A (i.e. on sessions 1 and 3), the viscosity coefficients on the right and left hands (*b*_*R*_, *b*_*L*_) were kept constant on all trials at (10,10) Ns/m. During the second session with task B, variability was introduced through the haptic forces. To introduce variability along the task-space, the viscosity coefficients were chosen so that they were positively correlated – for e.g., when the viscosity coefficient of the right hand for a trial was increased, the viscosity coefficient of the left hand for that trial was also increased so that both hands slowed down (relative to an unperturbed trial) during that trial. Since the speed of the puck was determined by the sum of the two hand speeds, a positive correlation in the viscosity coefficients tended to cause a change in the overall speed of the puck. On the other hand, to introduce variability along the null-space, the viscosity coefficients were negatively correlated – for e.g., when the viscosity coefficient of the right hand for a trial was increased, the viscosity coefficient of the left hand for that trial was decreased so that the right hand slowed down and the left hand sped up (relative to an unperturbed trial). This tended to maintain the speed of the puck close to the original value (since both hands usually tended to move with similar speeds). In addition to manipulating the space in which variability was introduced (task or null space), the magnitude of variability introduced (high or low) was determined by the magnitude of the change in the viscosity coefficients.

### Groups

Participants were randomly assigned to 4 groups (n =12 participants/group) in a 2 x 2 design where we manipulated the space in which variability was introduced (task or null space) and the amplitude of the variability introduced (high or low) (**Figure 1C**). For each group, the coefficients (*b*_*R*_, *b*_*L*_) were chosen on each trial from one of the following combinations (all units specified in Ns/m): (i) low task space variability group (AT’A) - (7;7), (10;10), (13;13), (ii) high task space variability group (AT”A) - (4;4), (10;10), (16;16), (iii) low null space variability group (AN’A) - (7;13), (10;10), (13;7), and (iv) high null space variability group (AN”A) - (4;16), (10;10), (16;4) Ns/m.

We also added two control groups (n = 12 participants/group). In the rest group (A0A), participants simply skipped the second session with no practice. In the no-variability group (AAA), participants performed the same task as the original task A with no haptic perturbations in the second session (i.e. with constant viscosity coefficients on both hands).

## Data analysis

Kinematic data recorded by the robot were sampled with a sampling frequency of 1 kHz. For every trial, we computed the speed of the two hands at the instant of release since this was the only determinant of performance in the task.

### Absolute Error

The absolute error was measured as the absolute difference between the speed of the puck and the desired speed (1.5 m/s). Lower absolute errors indicated better task performance.

### Task and null space variability

Due to the redundancy in the task, we partitioned movement variability in the hand speeds into a task and null space (Cardis et al. 2018). Given that the solution manifold in this task is a line (i.e. V_L_+V_R_ = 1.5 m/s), the variability along this manifold is the null space variability since it does not influence directly task performance, and the variability along the orthogonal direction (i.e. along the line V_L_ = V_R_) is the task space variability that influences task performance.

### Coordination strategy

Since the task was a precision task, we computed a coordination index that quantified the coordination strategy based on the relative amounts of task and null space variance in this task (Zhang et al. 2006). This was computed as:

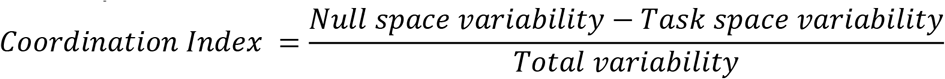

The value of the coordination index could range from −1 to +1. A positive value of coordination index indicates preferential use of the null space, whereas a negative value of coordination index indicates preferential use of the task space. Note that a separate normalization of the task and null space variability was not required in our case because both the task and null space had a dimension of 1.

## Statistical Analysis

### Effect of haptic perturbations on immediate performance

As a manipulation check, we first examined if the haptic perturbations had the desired effect on the task and null space variability when the perturbations were first introduced. We compared this using a 2 x 2 ANCOVA (Space x Amplitude) on the task and null space variability during the first block of task B (i.e., block 1 of Session 2) using the last block of practice of task A on the first day (i.e. block 8 of session 1) as the covariate. We repeated the same analysis on the absolute error and coordination strategy.

### Effect on haptic perturbations on motor memory consolidation

To examine consolidation, we used a 2 x 2 (Space x Amplitude) ANCOVA on the absolute error and the coordination index in task A after 24 hr (i.e. block 1 of session 3), using block 8 of session 1 as the covariate.

### Comparisons with the control groups

Finally, we compared each of the 4 groups against the control groups (A0A and AAA) by using two separate ANCOVAs for the absolute error and coordination index. Similar to the analysis of consolidation, this was conducted on the block 1 of session 3 using block 8 of session 1 as the covariate. A priori contrasts in each ANCOVA were restricted to comparison of the 4 groups against the control group (i.e. either A0A or AAA). In all cases, the significance level was set at .05.

## Results

### Outliers

Because variability measures are sensitive to the presence of outliers, we used the absolute error to filter out any outlier trials in each block. In each block, a trial was considered an outlier if the absolute error was outside the Tukey’s fences (i.e. less than Q1-1.5*IQR or greater than Q3+1.5*IQR, where Q1 and Q3 refer to the first and third quartiles, and IQR refers to the interquartile range). The total number of trials discarded was ~ 2%.

### Effect of haptic perturbations on immediate performance

We found that the haptic perturbations had the desired effect in the task and null space variability in the corresponding groups. For the task space variability, there was a significant effect of space F(1,43) = 7.739, p = .008, and amplitude F(1,43) = 9.594, p = .003 (**Figure 3A**). As expected, task space groups had higher task space variability than the null space groups, and high amplitude groups had higher task space variability than low amplitude groups. The space x amplitude interaction was not significant F(1,43) = .587, p = .448.

**Figure 3.**
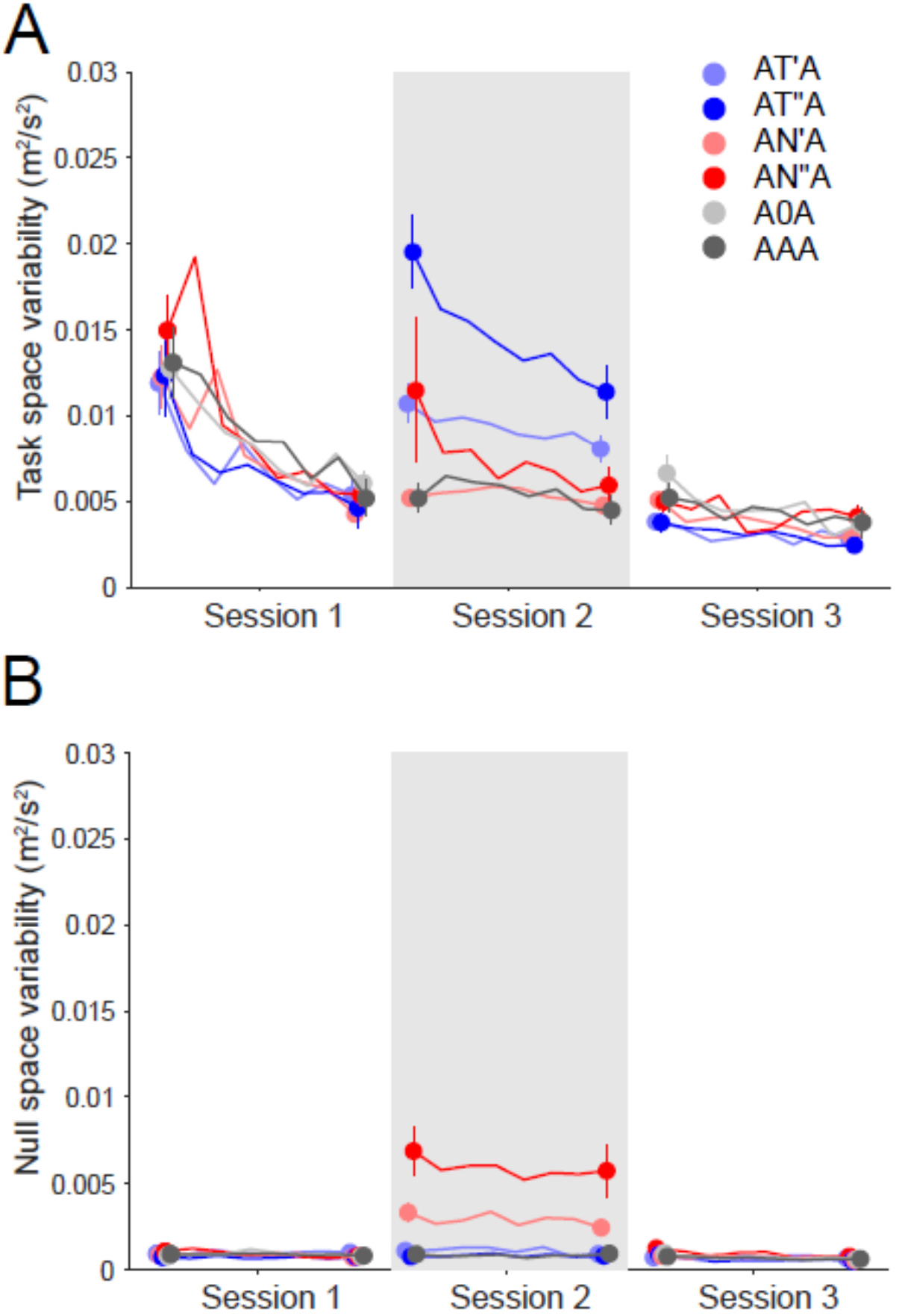
(A) Task and (B) null space variability during practice. During session 2 (when the haptic perturbations were introduced), task space variability was higher for the task space groups, and null space variability was higher for the null space groups. There was also an effect of amplitude with the high variability groups having more variability than the low variability groups.

For the null space variability, there were similar results – there was a significant effect of space F(1,43) = 31.83, p < .001 and amplitude F(1,43) = 4.97, p = .031 (**Figure 3B**). Again, as expected, null space groups had higher null space variability than the task space groups, and high amplitude groups had higher null space variability than the low amplitude groups. In addition, there was a Space x Amplitude interaction (F(1,43) = 5.42, p = .025) which indicated that for the null space groups, there was a significant difference in null space variability between the low and high amplitudes, whereas this was not the case for the task space groups.

For the absolute error, there was a main effect of space, F(1,43) = 21.09, p < .001 and amplitude, F (1,43) = 15.79, p < .001 (**Figure 4A**). Task space groups had higher absolute error than the null space groups, and the high amplitude groups had greater absolute error than the low amplitude groups. The space x amplitude interaction was not significant, F(1,43) = 1.59, p = .214.

**Figure 4.**
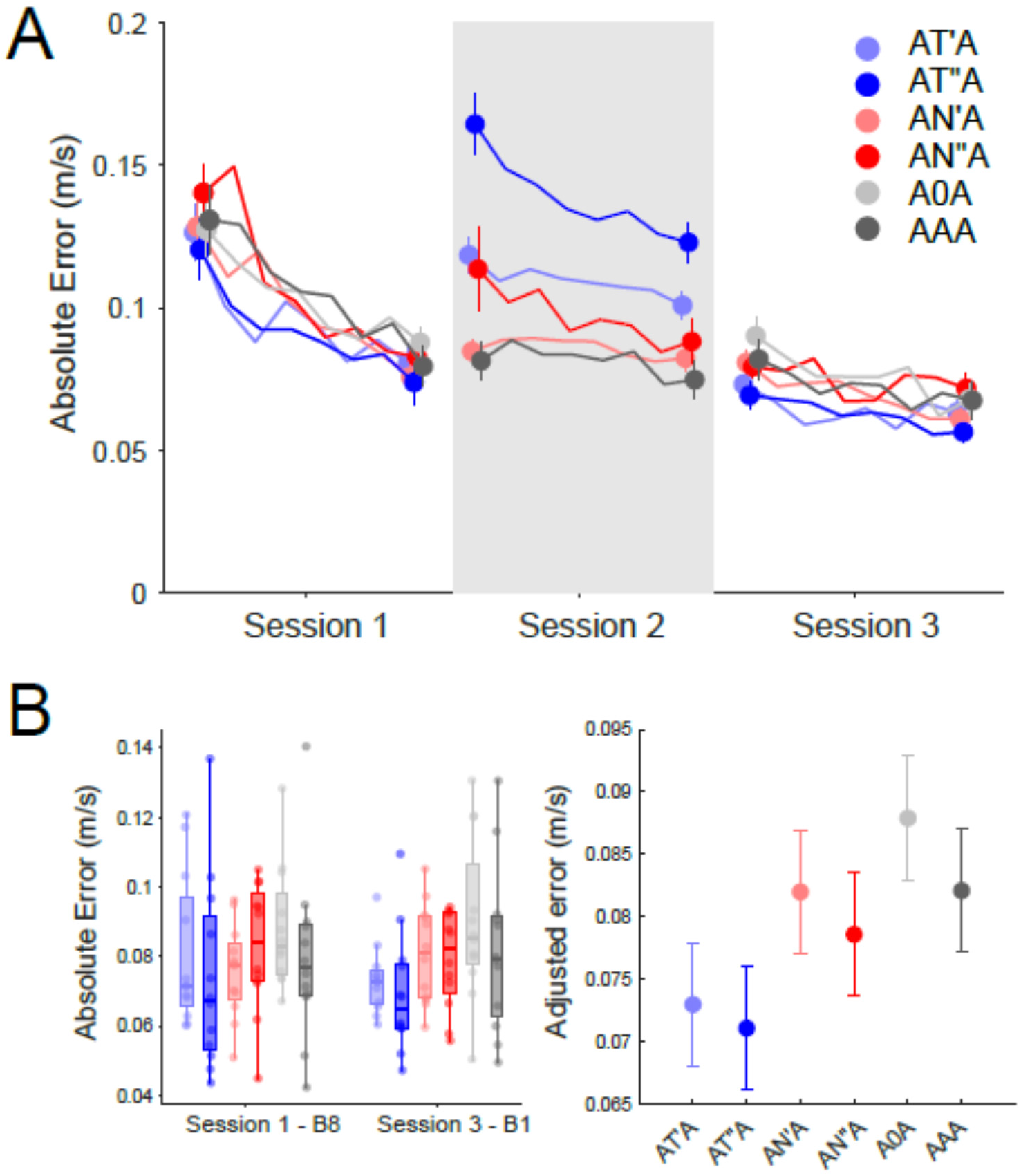
**(A) Absolute error during practice.** Practice resulted in reduced absolute error, although in session 2, the task space groups showed significantly higher error that the other groups. **(B) Consolidation effects.** Consolidation was analysed based on examining the error in the first block of session 3 relative to the last block of session 1 (using a covariate adjustment). Task space groups showed lower adjusted errors (i.e. better consolidation) relative to the null space groups and the A0A group.

For the coordination index, there was a significant main effect of space, F(1,43) = 71.509, p < .001. The null space groups had a significantly higher coordination index (i.e. relatively more variability distributed along the null space) relative to the task space groups (**Figure 5**). The main effect of amplitude, F(1,43) = .004, p =.945 and the space x amplitude interaction, F(1,43) = 1.465, p = .233 were not significant.

**Figure 5.**
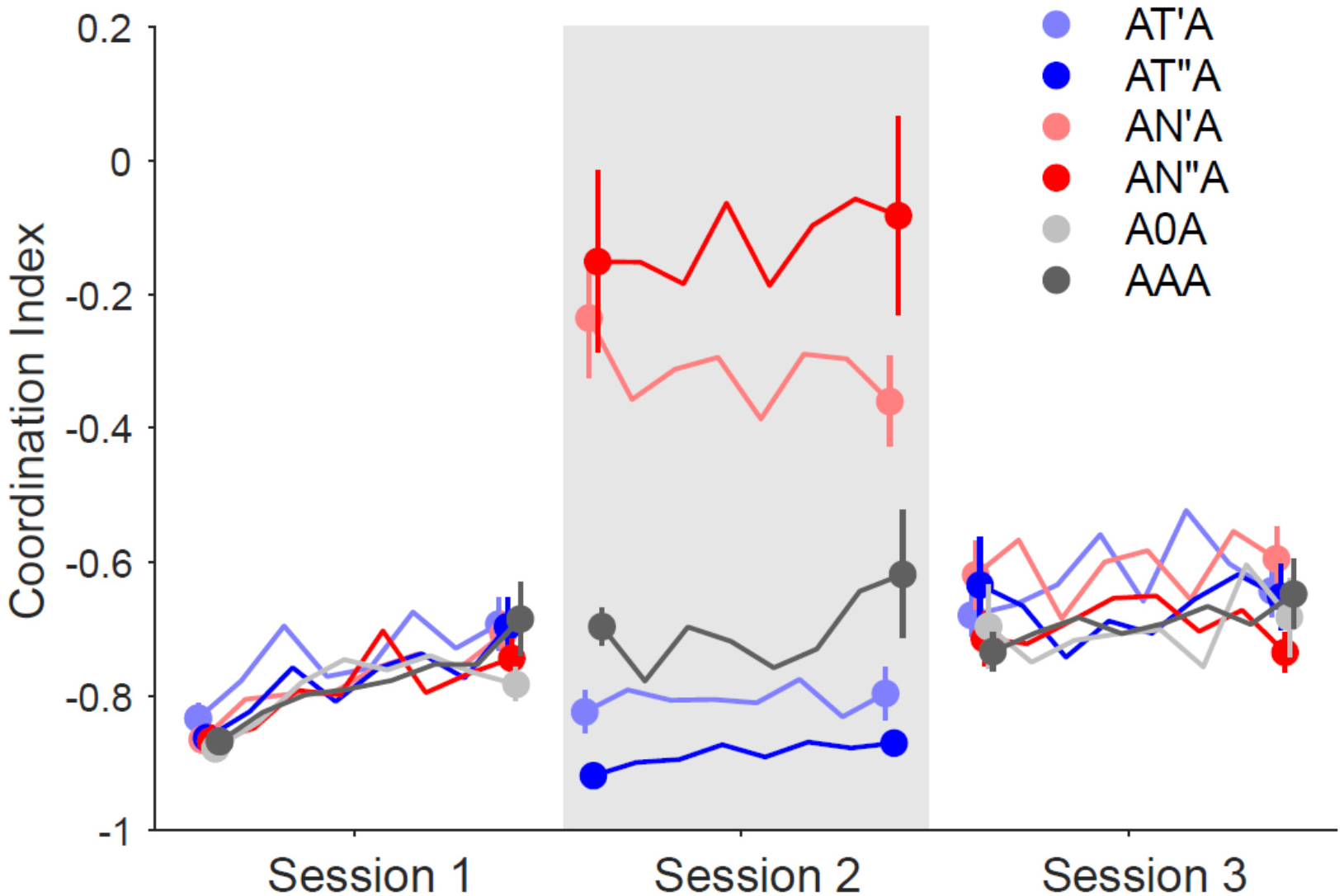
Coordination index during practice. Participants had a highly negative coordination index in session 1 (indicating higher variability in the task space). However, during session 2, the null space groups had a much higher coordination index (i.e. closer to zero) indicating that they were using a coordination pattern that was not congruent with that they had practiced in session 1. In session 3, all participants reverted to the highly negative coordination index once again, indicating that this was a preferred coordination strategy to perform the task.

### Effect of haptic perturbations on motor memory consolidation

For the absolute error, we found that task space groups had better consolidation relative to null space groups (**Figure 4B**). There was a significant main effect of space (F(1,43) = 4.561, p = .038), indicating that the task space groups had smaller absolute error (i.e. better consolidation) relative to the null space groups. The main effect of amplitude (F(1,43) = 0.467, p = .498) and the Amplitude x Space interaction (F(1,43) = .006, p = .939) were not significant.

For the coordination index, there were no statistically significant differences in the coordination strategy between groups after consolidation (**Figure 5**). The main effect of space F(1,43) = .002, p = .958, amplitude, F(1,43) = .123, p = .728, and space x amplitude interaction F(1,43) = 1.718, p = .197 were all not significant.

### Comparisons with control groups

#### Comparison with A0A

For the absolute error, we found that the task space groups had significantly lower absolute error (i.e. better consolidation) than the A0A group (p = .023 for ATA, p = .01 for AT”A). Both null space groups were not significantly different from the A0A group.

For the coordination index, there were no significant differences between any of the groups relative to A0A group.

#### Comparison with AAA

For both the absolute error and the coordination index, there were no significant differences between any of the groups relative to AAA (all ps > .05).

#### Exploratory analysis

To examine if the difference in consolidation between introducing variability in the task and null spaces lasted longer than just the initial block on Day 2, we conducted an exploratory analysis using a repeated measures ANCOVA across three blocks on Session 3 (Block 1, Block 4 and Block 8) with the block 8 of session 1 as covariate. We found a significant main effect of space (F(1,43) = 9.049, p =.004), indicating that the effects seen in the first block persisted throughout the rest of the learning (**Figure 4A**).

## Discussion

The goal of our study was to examine how the amplitude and space in which variability is introduced affect consolidation of a motor skill, which had redundant solutions. We found that the space in which variability was introduced had an effect on consolidation - variability in the task space resulted in better consolidation (i.e. lower errors on the 24 h period) compared to variability in the null space. In contrast, at least in the range studied, the amplitude of the variability had no noticeable effects on the consolidation. In addition, comparisons with control groups showed that: (i) both the task space groups performed better when compared to a control group that had no practice, but (ii) none of the groups were superior to intervening practice with no variability.

Why might introducing variability in the task space have a positive effect on consolidation? One potential explanation is that the variability in the task space created errors (as seen by the increase in absolute error in Session 2), which might have engaged error-based learning mechanisms, resulting in better consolidation. However, this explanation does not seem to fully account for the results as the amplitude of variability did not have an effect on the consolidation. For example, the errors experienced by the low task space variability group were lower than the high task space variability group - yet, in terms of consolidation observed, they were very similar.

A more likely explanation for the difference between the task and null space groups is suggested by the observation of how the imposed variability interacts with the pre-existing coordination strategy. During the practice of task A, participants generally used a coordination strategy with preferentially higher variability in the task space (i.e. a negative coordination index). Inducing variability in the task space did not disrupt this coordination strategy even though it increased the overall error. On the other hand, introducing variability in the null space significantly altered the coordination strategy and increased variability along the null space (making the coordination index almost close to zero). This meant that even though overall task error was not high, these groups were practicing a coordination strategy that was different from their preferred strategy to solve the task. The assumption that the pre-existing coordination strategy was a ‘preferred strategy’ in the task is also supported by the observation that on session 3 (24 h after the first session), the null space variability groups (which had practiced a different coordination strategy when the haptic perturbations were present) spontaneously switched back to this strategy.

These results are consistent with a view that learning does not occur on a ‘blank slate’, but instead builds on pre-existing coordination tendencies (Kostrubiec and Zanone 2002; Zanone and Kelso 1992). The tendency to distribute variability along the task space (even though it is suboptimal from a task performance standpoint) is consistent with other prior work showing that the two hands tend to be coordinated in a ‘symmetrical’ pattern (Kelso et al. 1979; MacKenzie and Marteniuk 1985) – i.e., the tendency to specify the same movement parameters to both limbs when they are moved simultaneously (Swinnen and Wenderoth 2004). When this preferred coordination pattern is interfered with (in this case by increasing the null space variability), subsequent consolidation is affected, even though immediate task performance itself was not. Moreover, our exploratory analysis indicated that these differences not only affected the consolidation in terms of the performance immediately after the break, but also had rather prolonged effects on learning, lasting at least for another 400 trials of practice. These results support the idea that the influence of variability on learning is not merely restricted to its effect on task performance, but also on how the imposed variability interacts with the prior coordination patterns (Ranganathan and Newell 2013).

Our two control comparisons (with A0A and AAA) indicated that the introduction of task space variability had better consolidation relative to the A0A group but not the AAA group. Although the task space groups tended on average to have better consolidation even relative to the AAA group, these changes were not significant, and therefore it is not clear if there is a benefit to introducing task space variability relative to no variability. Given the relatively small sample size used, a more high-powered replication may be necessary to tease out this effect.

Interestingly, we found that the amplitude of variability introduced (at least in the range here) did not have an influence on consolidation. In a prior study using the same task with the same range of variation (Cardis et al. 2018), we found that higher amplitudes of variability was detrimental to learning when the variation was introduced during the practice of the original skill, instead of as an interfering task as done here. These results suggest that the influence of variability on learning may depend on the time course of the learning – i.e. the amount of variability may be critical in influencing learning during initial acquisition of the task, but may not be as critical once that memory has consolidated.

There are two important distinctions from prior work that are worth mentioning. First, the issue of how variability is introduced is an important consideration for consolidation and learning. Wymbs and colleagues (Wymbs et al. 2016) make the distinction between endogenous (intrinsic) and exogenous (externally induced) variability. However, even within exogenous variability, there may be differences in how variability is introduced. In the Wymbs et al. study, the variability in the task was introduced through a change in the visuomotor mapping, which meant that there were no direct perturbations to the movement and participants had to produce variability on their own to match the task goal. However, in our case, variability was introduced directly through mechanical perturbations of the movement. The use of mechanical perturbations was preferred in our context because while variability in the task space can be created through change in visual feedback, there is no direct way of increasing null space variability without introducing a change to the task in some way (for example by introducing a secondary constraint). Although these perturbations were introduced smoothly, since they were caused by a change in the viscosity coefficients that are proportional to the velocity (and not by a discrete ‘pulse’ of force), the results suggest that even within exogenous variability, self-generated and externally-imposed variability may have different effects on learning and consolidation. A second deviation from the literature, as mentioned in the introduction, relates to the nature of the task and the use of redundancy. Much of prior work on consolidation examines the effects of adaptation or sequence learning (Brashers-Krug et al. 1996; Krakauer and Shadmehr 2006; Robertson et al. 2004; Wymbs et al. 2016). Learning to control variability, on the other hand, has been much less investigated (Muller and Sternad 2004; Ranganathan and Newell 2010; Sternad et al. 2011) and likely engages different learning mechanisms from these other tasks (Krakauer et al. 2019; Shmuelof et al. 2012, 2014). In particular, the presence of redundancy in the task allows a closer look at how consolidation impacts learning beyond the level of task performance and into the level of coordination strategies (since multiple coordination strategies can result in the same task performance). Combined with other work in the study of such tasks (Levac et al. 2019; Sternad 2018), we suggest that the study of redundancy may provide further insight into how learning and consolidation are affected in real-world tasks.

In summary, we found that the space in which variability was introduced had distinct effects on consolidation of motor memories. These results suggest that the effects of variability on motor memory consolidation depend on the interplay between the imposed variability and the pre-existing coordination strategy and highlight the need to consider coordination as a critical element in motor memory consolidation of tasks with redundancy.

## Acknowledgment

This work was supported by NSF grant 1823889. MP was partially supported by the operative program Por FSE Regione Liguria 2014-2020 RLOF18ASSRIC/15/1.

